# Tumor secreted extracellular vesicles regulate T-cell costimulation and can be manipulated to induce tumor specific T-cell responses

**DOI:** 10.1101/2020.04.22.055632

**Authors:** Xianda Zhao, Ce Yuan, Dechen Wangmo, Subbaya Subramanian

## Abstract

Tumor intrinsic factors negatively regulate tumor immune cell infiltration and function. Deciphering the underlying mechanisms is critical to improving immunotherapy in cancers. Our analyses of human colorectal cancer (CRC) immune profiles and tumor-immune cell interactions revealed that tumor cell secreted extracellular vesicles (TEVs) induced immunosuppression in CRC. Specifically, TEVs containing microRNA miR-424 suppressed the CD28-CD80/86 costimulatory pathway in tumor infiltrating T cells and dendritic cells. Modified TEVs with miR-424 knocked down enhanced T-cell mediated antitumor immune response in CRC tumor models and increased the response to immune checkpoint blockade therapies (ICBT). Intravenous injections of modified TEVs induced tumor antigen specific immune responses. Moreover, injections of modified TEVs boosted the ICBT efficacy in CRC models that mimic treatment refractory late-stage disease. Collectively, we demonstrate a critical role for TEVs in antitumor immune regulation and immunotherapy response, which could be developed as a novel treatment for ICBT resistant human CRC.

## INTRODUCTION

Therapies targeting immune checkpoint pathways including the cytotoxic T lymphocyte-associated protein (CTLA-4) and the programmed cell death-1 (PD-1)/PD-1 ligand-1 (PD-L1), have revolutionized cancer treatment(Darvin et al., 2018; Hargadon et al., 2018; Sharma and Allison, 2020). For colorectal cancer (CRC), immune checkpoint blockade therapy (ICBT) is effective in 30-60% of the microsatellite instable-High (MSI-H) subtype(Oliveira et al., 2019). Unfortunately, most CRC patients (>85%) have microsatellite stable (MSS) tumors(Kim et al., 2016; Mlecnik et al., 2016), that do not respond to ICBT. The MSI-H tumors are generally associated with a higher mutational load and are considered immunogenic(Salem et al., 2018; Zhao et al., 2018). Currently, the MSI/MSS phenotype is considered as a biomarker for ICBT response(Le et al., 2017; Mandal et al., 2019). However, this MSI/MSS based stratification alone cannot adequately explain the observed difference in treatment response since immune cell infiltration is also observed in a large proportion of MSS tumors(Mlecnik et al., 2016; Pages et al., 2018). A mechanistic understanding of why the existing tumor infiltrating immune cells are not functionally rescued by ICBT is critical for improving outcomes in both MSS- and MSI-CRC patients.

Intercellular communications between tumor cells and immune cells in the tumor microenvironment (TME) affect the antitumor immune response and the tumor response to ICBT. One way tumor cells can affect surrounding cells is by secreting extracellular vesicles (EVs)(Kalluri and LeBleu, 2020) and tumor cell-derived EVs (TEVs) can be either immunogenic or immunosuppressive depending on their cargo. For example, PD-L1 in TEVs have strong immunosuppressive effects(Chen et al., 2018; Poggio et al., 2019), while the tumor antigens and other immune stimulating factors carried in EVs can induce immune responses(Diamond et al., 2018; Horrevorts et al., 2019; Wolfers et al., 2001). Investigation of additional tumor intrinsic mechanisms and immunosuppressive components in EVs that incite ICBT failures are highly warranted.

It has been demonstrated that CD28-CD80/86 costimulatory signals are critical for ICBT response(Hui et al., 2017; Kamphorst et al., 2017). For example, PD-1 recruits the Shp2 phosphatase, which preferentially inhibits CD28 signaling(Hui et al., 2017). This suggests that anti-PD-1 efficacy is, in part, due to rescuing CD28-CD80/86 signaling. Another study demonstrated that CD28-CD80/86 signaling in tumor infiltrating CD8^+^ T cells is essential for anti-PD-1 response(Kamphorst et al., 2017). Markedly, CD80 expression on the antigen presenting cells (APCs) was recognized as a novel mechanism that restricts PD-1 signaling on interacting T cells by forming cis-heterodimers with PD-L1(Sugiura et al., 2019). This CD80-PD-L1 interaction inhibits CD80 signaling through CTLA-4 thus attenuating the immunosuppression mediated by CTLA-4(Zhao et al., 2019). These reports led us to ask if other mechanisms, particularly tumor-intrinsic ones, led to CD28-CD80/86 signal dysregulation. We found this signaling axis was disrupted in human CRC patients by TEVs. Additionally, we investigated if depleting the potential CD28-CD80/86 suppressive factors in TEVs would enhance their immunogenic effects, thus improving antitumor immunity.

## RESULTS

### Costimulatory molecules CD28, CD80, and CD86 are downregulated on human CRC infiltrating immune cells

We determined the overall adaptive antitumor immune response in human CRC tissues (24 MSI subtype and 47 MSS subtype) by examining the T-cell infiltration in the TME. Tumors with well- and poorly-infiltrated T cells were observed in both microsatellite instable (MSI) and microsatellite stable (MSS) subtypes (Figure 1A). Overall, the MSI-CRC had significantly higher numbers of tumor infiltrating T cells compared to MSS-CRC (Figure 1A). This observation corroborated our immune signature analysis of the TCGA-CRC dataset (Figure S1A) and previous reports in CRC(Becht et al., 2016; Llosa et al., 2015; Mlecnik et al., 2016).

**Figure 1.**
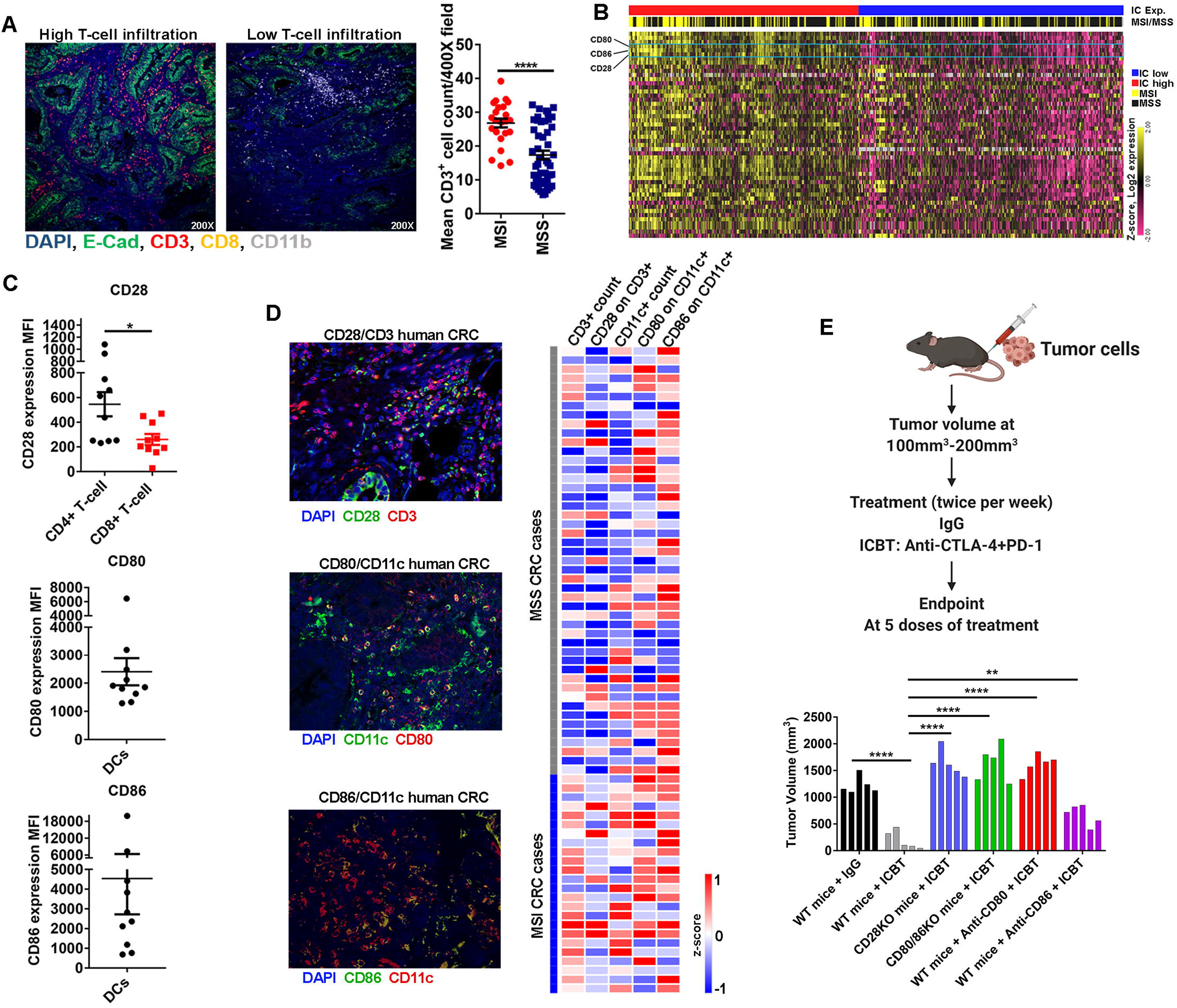
Expression of costimulatory molecules CD28, CD80, and CD86 on tumor infiltrating immune cells in human CRC. (A) Multiplex immunofluorescence staining of tumor infiltrating T cells (CD3, CD8) and myeloid cells (CD11b) in human CRC tissue samples. (24 MSI cases, 47 MSS cases, Mean±SEM, t-test, \*\*\*\**p*<0.0001) (B) Immune checkpoints expression profile in TCGA CRC dataset. (C) Flow cytometry analysis of CD28, CD80 and CD86 on immune cells in fresh human CRC tissues. (Mean±SEM, n=10, t-test, \**p*<0.05) (D) Multiplex immunofluorescence staining of CD28 on T cells and CD80/86 on DCs in a human CRC cohort (24 MSI cases, 47 MSS cases). Only T cells and DCs in the tumor center were scored. Magnification 400X. (E) Impact of CD28 and CD80/86 expression on ICBT (anti-CTLA-4 + anti-PD-1) in subcutaneous tumors. ICBT restricted early stage tumor development in WT mice. However, *Cd28*^*-/-*^ mice and *Cd80/86*^*-/-*^ mice were resistant to ICBT. Antibody blocking of CD80 and CD86 induced tumor resistant to ICBT. (Individual data for each case, n=5, t-test, \*\**p*<0.01, \*\*\*\**p*<0.0001) See also Figure S1.

Next, we measured the immune checkpoint genes (ICG) expression in CRC. Genes expression analysis of a set of 50 ICGs, including CD28, CD80, and CD86, in TCGA-CRC dataset stratified tumors into ICG-high and -low expression groups. Both MSI- and MSS-CRC tumors were seen in each group (Figure 1B). At the protein level, we measured CD28-CD80/86 costimulatory molecules on human CRC infiltrating T cells and dendritic cells (DCs). Flow cytometry analysis of a cohort of 10 primary human CRC fresh tissues showed that CD4^+^ T cells generally had higher levels of CD28 than CD8^+^ T cells (Figures 1C & S1B). However, variable levels of CD28 expression were observed on both CD4^+^ and CD8^+^ T cells (Figure 1C). Likewise, tumor infiltrating DCs had variable levels of CD80/86 (Figure 1C). These data, though, are composites of TILs and DCs throughout the entirety of the tumor, and do not account for potential spatial differences. To specifically study the CD28-CD80/86 costimulatory molecules on T cells and DCs localized in the center of tumors, we collected 71 archived human CRC tissues (24 MSI subtype and 47 MSS subtype) for immunofluorescence analysis. Only T cells and DCs that intercalated within or between the epithelial component of the CRC/normal colon tissue were selected for scoring (Figures 1D & S1C-D). No/low expression of CD28 (score<0.5, Figure S1C) on T cells was observed in ∼60% of all cases in both MSI- and MSS-CRC tumors (Figure 1D). Similarly, DCs located in the central tumor showed no/low expression of CD80/86 (score<0.5, Figure S1C) in ∼50%/∼33% of all cases (cutoff=0.5, Figure 1D). Taken together, our data showed that the expression of CD28 and CD80/86 is highly variable and can be weak or absent on tumor infiltrating T cells and DCs in both MSI- and MSS-CRC tumors.

### CD28-CD80/86 costimulatory pathway is necessary for ICBT response

Having observed weak/no expression of CD28-CD80/86 on tumor infiltrating immune cells in human CRC, we tested whether loss of CD28-CD80/86 expression affected tumor response to ICBT in a subcutaneous mouse model of CRC. Our data showed that a combination of anti-PD-1 and anti-CTLA-4 could significantly suppress growth of early stage (100-200mm^3^) subcutaneous tumors (MC38 mouse CRC cell line with MSI phenotype) (Figure 1E). Single drug treatment with either anti-PD-1 or anti-CTLA-4 was found to be less effective than combination treatments (data not shown). However, in both *Cd28*^*-/-*^ and *Cd80/86*^*-/-*^ mice, the combination treatment did not suppress growth (Figure 1E & S1E). In human CRC, loss of CD28 and CD80/86 expression occurs in response to a tumor-dependent mechanism, rather than a genetic deficient mechanism. To model this, we administered anti-CD80 and anti-CD86 blocking antibodies to wild type mice bearing early stage tumors to mimic acquired deficiency (depletion after tumor growth). Notably, blocking CD80 after the initial tumor development abolished the efficacy of ICBT (Figure 1E & S1E). Blocking CD86 showed a similar but weaker effect than the CD80 (Figure 1E & S1E). Reanalysis of published data(Prat et al., 2017) indicated that tumors responding to immune checkpoint inhibitors have higher levels of CD28 and CD80/86 expression (Figure S1F). These results suggested that adequate CD28 and CD80/86 expression were necessary for effective ICBT and these findings are consistent with previous studies(Hui et al., 2017; Kamphorst et al., 2017).

### MicroRNA-424 is overexpressed in human CRC and inhibits CD28 and CD80 expression

MicroRNAs (miRNAs) are small non-coding RNAs that post-transcriptionally regulate gene expression(Bracken et al., 2016). To determine if miRNAs regulate CD28-CD80/86 costimulatory molecules in human CRC, we first identified miRNAs that target CD28 and CD80/86 by target prediction (Figure 2A). TCGA-CRC data revealed that miR-424 is upregulated in CRC but downregulated in normal colon tissues (Figure 2A-B). By *in situ* hybridization (ISH), we validated that miR-424 is expressed in cancer tissue, but not in normal colon tissue (Figure 2C and Figure S2A). In another cohort of 21 human CRC and patient-matched normal tissues, we quantified miR-424 expression by qRT-PCR (Figure 2D). Fourteen out of 21 CRC cases showed at least 2-fold upregulation of miR-424 in contrast to corresponding normal colon tissues (Figure 2D). Hence, miR-424, that potentially inhibits CD28 and CD80 expression appears upregulated in human CRC. We next sought to confirm the inhibitory effects of miR-424 on CD28 and CD80. In dual luciferase reporter vectors, we inserted the 3’UTR sequences of CD28 or CD80 (Figures 2E and S2B).

**Figure 2.**
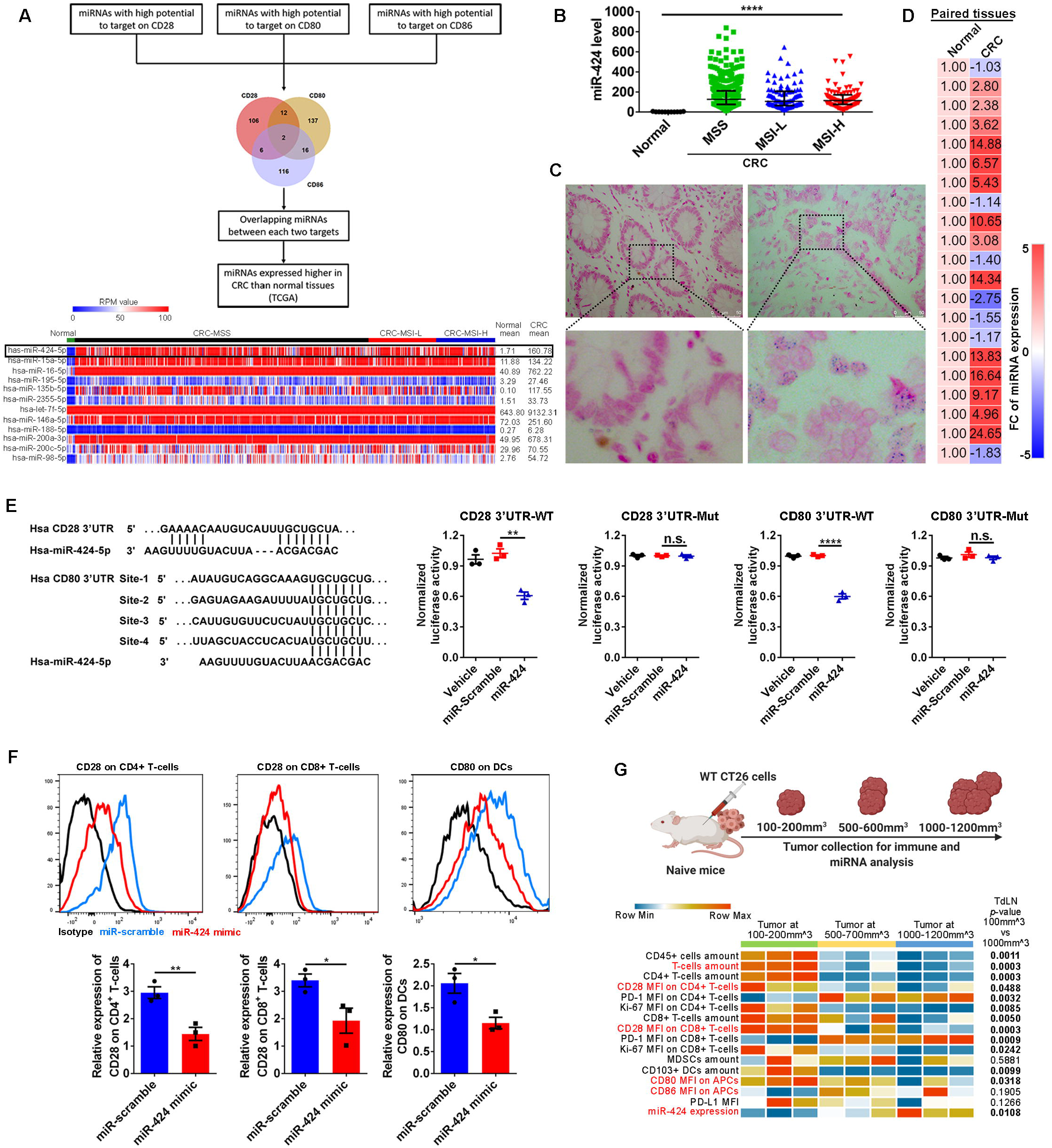
miR-424 is highly expressed in human CRC and negatively regulate CD28 and CD80 expression. (A) Schematic of process to identify miR-424 as a mechanism to downregulate the CD28 costimulatory pathway in human CRC. (B) miR-424 expression in human CRC and normal colon cases (TCGA dataset). (Kruskal-Wallis test, \*\*\*\**p*<0.0001) (C) In situ hybridization (ISH) of miR-424 expression in human tissues. Representative staining of normal colon (left panel) and CRC tissue (right panel). (n=10) (D) qRT-PCR analysis of miR-424 expression in paired human CRC and normal colon tissues. Scale shows Fold change (FC). (E) Dual luciferase reporter assay of miR-424 on the 3’UTR of human CD28 and CD80. (Mean±SEM, n=3, t-test, \*\**p*<0.01, \*\*\*\**p*<0.0001, n.s. no significance) (F) Primary human CD4^+^ and CD8^+^ T cells, and DCs transfected with miR-424 mimic showed lower CD28 and CD80 protein expression respectively. (Mean±SEM, n=3, t-test, \**p*<0.05, \*\**p*<0.01) (G) Heatmap representation of miR-424 expression level and the immune landscapes during various phases of CT26 tumor growth. (n=3, t-test, *p*-values listed) See also Figure S2.

Reporter assays showed that miR-424 significantly reduced the luciferase signal (Figures 2E and S2B). In the reciprocal experiments, site-directed mutations on the 3’UTRs nullified the effects of miR-424 (Figure 2E, Figure S2B). We then isolated primary human T cells and DCs that endogenously expressed CD28 or CD80 from healthy donors’ blood and transfected them with miR-424 or scrambled miRs (Figure 2F). Again, miR-424 showed a strong inhibitory effect on CD28 and CD80 protein expression (Figure 2F). These data confirm that miR-424 binds to 3’UTRs of CD28 and CD80 and suppress their expression on T cells and DCs, respectively.

Subsequently, we extended our investigations to mouse CRC models established by subcutaneous injections of CT26 (MSS phenotype) and MC38 (MSI phenotype) mouse CRC cell lines. We assessed the immune cell landscape and miR-424 expression changes during different stages of tumor growth (100-200mm^3^, 500-700mm^3^, and 1000-1200mm^3^). Notably, we observed a gradient decrease of tumor infiltrating immune cell frequencies and CD28 and CD80/86 expression during tumor development (Figures 2G and S2C-E). In contrast, miR-424 expression significantly increased with tumor growth in both CT26 and MC38 tumors (Figures 2G and S2C-E). These findings indicate that increased miR-424 expression is correlated with low CD28 and CD80 protein expression and an immunosuppressive phenotype during tumor growth.

### TEVs transfer miR-424 from human CRC cells to tumor infiltrating T cells and DCs

As our data indicated that miR-424 upregulation in human CRC was a potent inhibitor of CD28 and CD80 expression, we sought to identify the mechanism that led to the accumulation of miR-424 in human CRC and tumor infiltrating T cells and DCs. Strikingly, the level of miR-424 was significantly higher in human and mouse CRC cell lines and primary human CRC tumor organoids, than to the species matched naïve T cells and DCs (Figure 3A-B). Extracellular vesicles (EVs) are critical in mediating the transfer of microRNAs and intercellular communication in the TME(Huber et al., 2018; Kalluri and LeBleu, 2020; Valadi et al., 2007). Therefore, we investigated whether tumor cell expressed miR-424 could be transferred to T cells and DCs, leading to strong functional effects in the recipient cells. TEVs enriched for exosomes (average size ∼130nm) from human and mouse CRC cell lines were isolated and characterized (Figure S3A-C). In these TEVs, we observed high levels of miR-424 by absolute qRT-PCR (Figure 3A). Using confocal microscopy, we confirmed uptake of fluorescence labeled TEVs by T cells and DCs cells *in vitro* (Figure 3C). We then examined TEVs mediated miR-424 transfer, in a transwell coculture system that contained tumor cells and T cells or DCs. Notably, the presence of tumor cells significantly increased the miR-424 level in T cells and DCs (Figure 3D). Treatment with the sphingomylenase inhibitor GW4869 could block TEVs production without affecting tumor cell viability (Figure S3D). When TEVs production is blocked by GW4869, tumor cells failed to cause an increase in miR-424 levels in cocultured T cells and DCs (Figure 3D).

**Figure 3.**
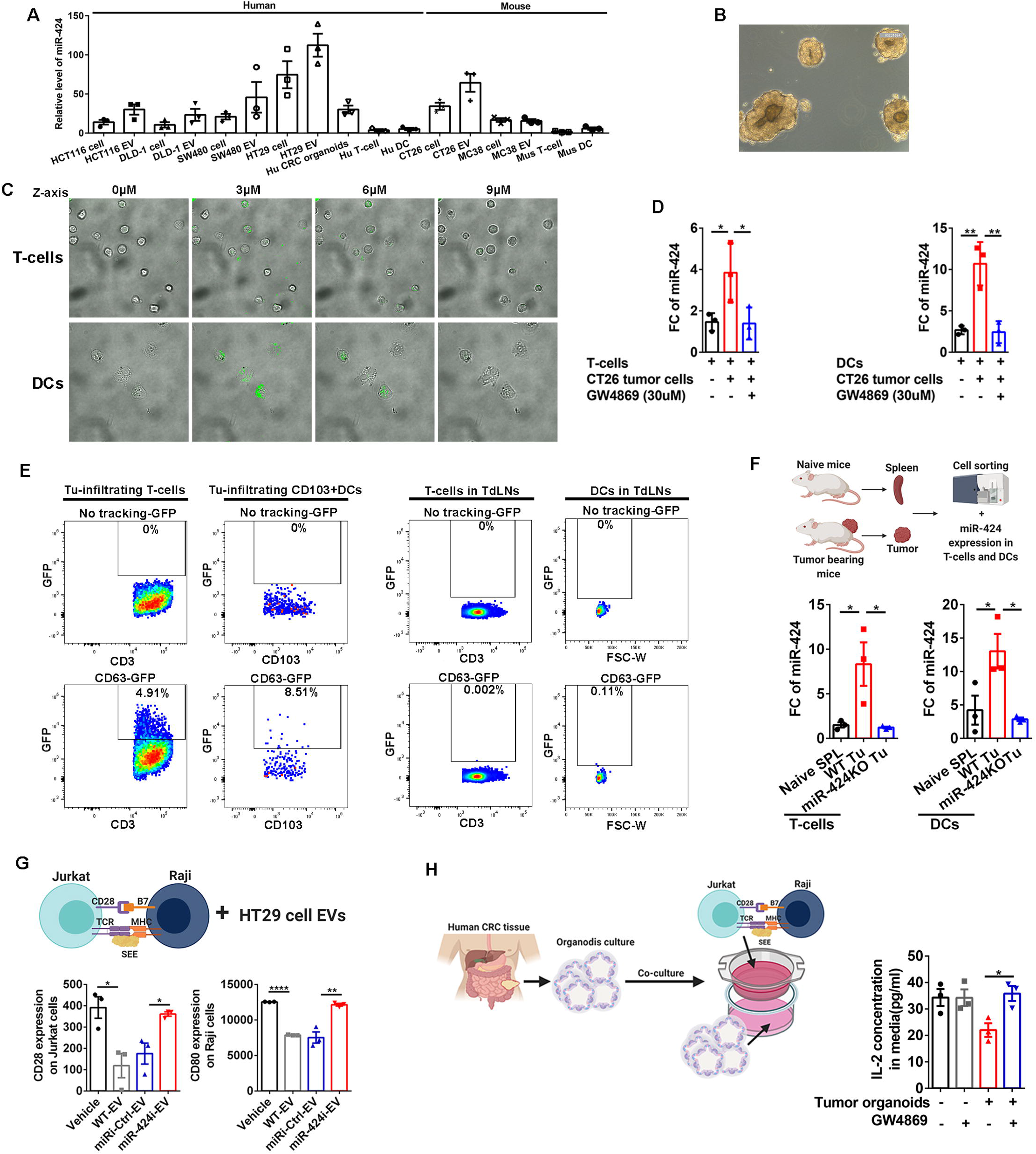
Tumor cells derived EVs transfer miR-424 to immune cells and downregulate CD28 and CD80. (A) Relative levels of miR-424 expression in human and mouse CRC cell lines, primary human and mouse T cells and DCs, and primary human CRC organoids. (B) Representative photomicrograph of primary human CRC organoids. (C) Confocal microscopy confirmed EVs up take by T cells and DCs. CT26 cells derived EVs were stained with Dio dye and incubated with mouse naïve T cells and DCs. (D) CT26 cells cocultured with naïve T cells and DCs in a transwell system with and without EV inhibitor GW4869. miR-424 levels in T cells and DCs in each experimental condition. (Mean±SEM, n=3, t-test, \**p*<0.05, \*\**p*<0.01) (E) Flow cytometry analysis showed GFP signal in a proportion of tumor infiltrating T cells and DCs of the CT26-CD63-GFP tumors. Wild type (upper row) and CD63-GFP transfected CT26 cells (lower row) were used to establish subcutaneous tumors. T cells and DCs in the CT26-CD63-GFP tumor draining lymph nodes (TdLNs) showed very weak GFP signal. (n=3 in each condition) (F) miR-424 levels in T cells and DCs isolated from the spleen of naïve mice and subcutaneous tumors (established by wild type (WT) tumor cells or miR-424 knockout (KO) tumor cells) by flow sorting. (Mean±SEM, n=3, t-test, \**p*<0.05) (G) Expression levels of CD28 on Jurkat cells and CD80 on Raji cells in conjugation system: WT-EVs, miRi-Ctrl-EV, and miR-424i-EV derived from human HT29 CRC cells were used for each set. (Mean±SEM, n=3, t-test, \**p*<0.05, \*\**p*<0.01, \*\*\**p*<0.001) (H) Human CRC derived organoids were cocultured with the Jurkat and Raji cells conjugation system in a transwell system with or without GW4869. IL-2 production in the coculture system was measured. (Mean±SEM, n=3, t-test, \**p*<0.05) See also Figure S3.

We extended our *in vitro* work to mouse models to validate that TEVs are responsible for transferring miR-424 from tumor cells to T cells and DCs in a physiologically relevant condition. WT CT26 cells or CT26 cells expressing CD63-GFP (a marker to trace EVs) were used to establish *in vivo* tumors. The GFP signal was detected in tumor (CT26-CD63-GFP) infiltrating T cells and DCs (Figure 3E). However, T cells and DCs in the tumor draining lymph nodes (TdLNs) and spleen showed no/weak GFP signal (Figures 3E and S3F). T cells and DCs sorted from mice implanted with WT tumors by flow cytometry showed higher levels of miR-424 than T cells and DCs isolated from mice implanted with miR-424 knockout (miR-424KO) tumors and naïve mice spleen (Figure 3F). Further, we determined that activation of T cells and DCs did not regulate their endogenous miR-424 expression (Figure S3G). Thus, the accumulation of miR-424 in tumor infiltrating T cells and DCs is due to the uptake of miR-424 containing TEVs.

Next, we determined the functionality of TEVs with/without functional miR-424 on the CD28-CD80/86 costimulatory pathway. We first characterized TEVs isolated from WT tumor cells, miR-424 inhibitor (miR-424 antisense oligonucleotides) transfected tumor cells, and miR-424KO tumor cells. We confirmed that tumor cells expressing the miR-424 inhibitor packaged the inhibitor into secreted EVs (miR-424i-EV) (Figure S3H). Further, we validated that miR-424i-EV and miR-424KO-EV (EVs derived from miR-424KO tumor cells) do not have functional miR-424 in them and cannot regulate CD28 and CD80 (Figure S3I). In a human *in vitro* model, we incubated Jurkat T cells with the Raji Burkitt’s lymphoma cell line to mimic a T cell-APC cell interaction. Addition of TEVs derived from the human CRC cell line HT29, which have normal miR-424 inhibited CD28 and CD80 expression on Jurkat and Raji cells, respectively (Figure 3G). Transfecting HT29 with a miR-424 knock-down construct abolished this effect, while transfection with a control did not (Figure 3G). Next, we tested the effect of TEVs generated from CRC patient derived tumor organoids on T-cell activation, which is indicated by IL-2 production. Tumor organoids retain the heterogeneity of tumors and are considered to be advanced *in vitro* models(Kopper et al., 2019; van de Wetering et al., 2015). In this experiment, we used a transwell assay with Jurkat/Raji cells in the upper chamber and CRC organoids in the lower chamber (Figure 3H). The presence of tumor organoids reduced IL-2 production from the Jurkat-Raji conjugation system. Noticeably, blocking EVs secretion by GW4869 rescued IL-2 production (Figure 3H).

Finally, we investigated how miR-424 expression is upregulated and regulated in human CRC cells. Hypoxia is a key feature that differentiates tumors from normal tissues and is commonly observed in human CRC(Baba et al., 2010). Induction of hypoxia stabilized the transcription factor HIF-1α expression in HT29 cells and led to the upregulation of miR-424 and increased EV production (Figure S3J). Our findings align with previous reports in other experimental systems(Ghosh et al., 2010; King et al., 2012; Zhang et al., 2014). Collectively, these data reveal that hypoxia in tumor cells causes upregulation of miR-424 that is delivered to tumor infiltrating T cells and DCs by TEVs where it suppressed CD28-CD80/86 costimulatory signaling.

### Blocking tumor cell derived miR-424 inhibits tumor development in an immune-dependent manner

Because TEVs with miR-424 suppressed the CD28-CD80/86 costimulatory pathway, we hypothesized that tumor-derived miR-424 impacts mouse CRC tumor development *in vivo*. Stably blocking the expression of Rab27a (a protein regulating exosome secretion) using shRAB27a reduced TEV production by MC38 tumor cells (Figure 4A). Suppressing TEVs’ secretion significantly reduced MC38 tumor growth *in vivo* (Figure 4B). In another experiment, mice were subcutaneously injected with tumor cells that cannot produce endogenous miR-424 containing TEVs. The resultant tumors were then injected with TEVs containing functional miR-424 which significantly increased tumor growth (Figure 4C). These data demonstrate that miR-424 in TEVs accelerates tumor development in immune competent mice.

**Figure 4.**
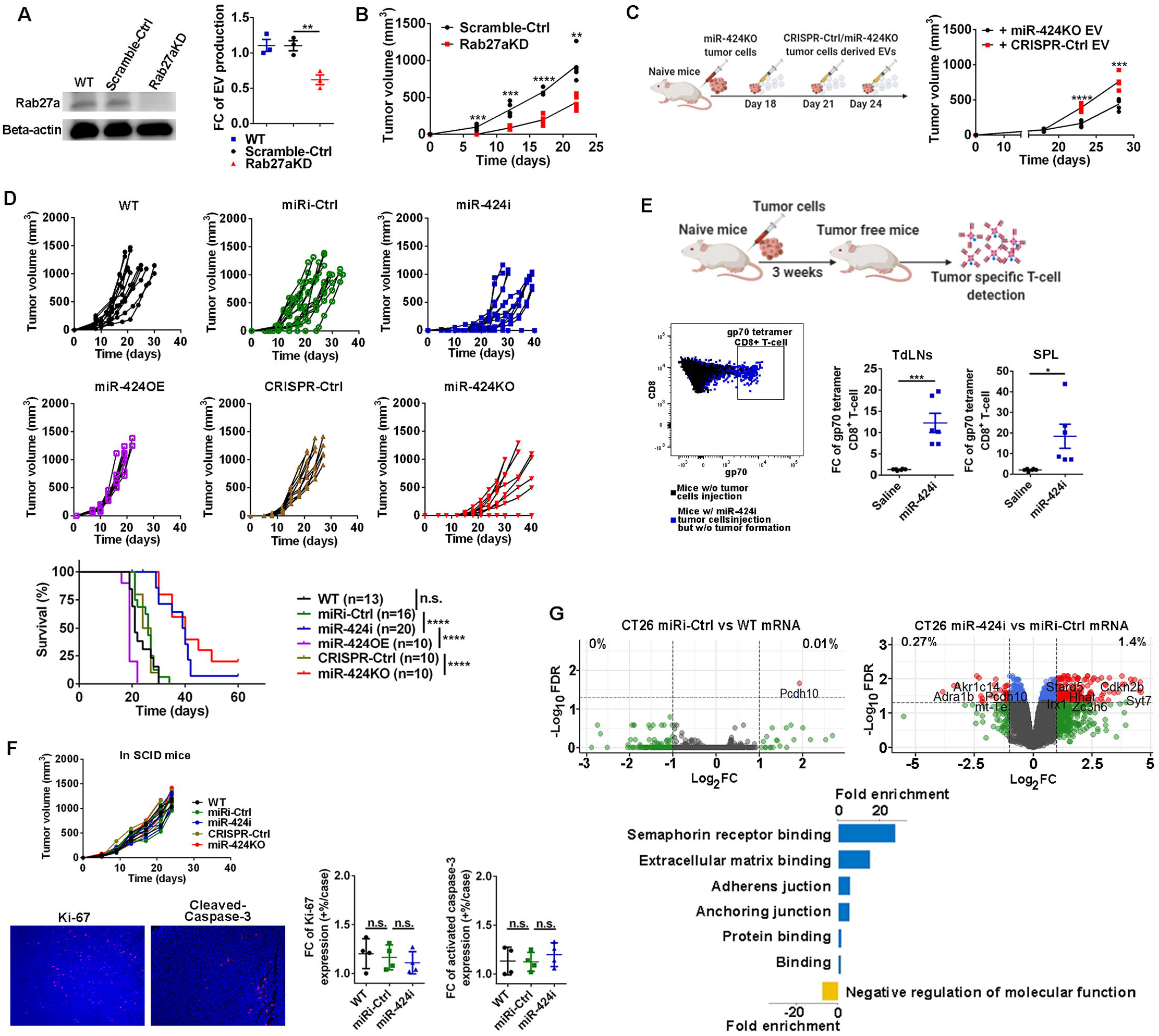
Tumor EVs with functional miR-424 accelerated tumor growth by suppressing antitumor immunity. (A) Knockdown of Rab27a significantly reduced EVs production in MC38 cells. (Mean±SEM, n=3, t-test, \**p*<0.05) (B) Growth curve of MC38 subcutaneous tumors established by MC38-Rab27aKD and MC38-Scramble-Ctrl cells. (Mean±SEM, n=5, t-test, \*\**p*<0.01, \*\*\**p*<0.001, \*\*\*\**p*<0.0001) (C) CT26 EVs with functional miR-424 (CRISPR-Ctrl EV) or without miR-424 (miR-424KO EV) were injected into the CT26-miR-424KO tumors in mice. Exogenous compensation of TEVs with miR-424 enhanced growth of tumors without endogenous miR-424. (Mean±SEM, n=5, t-test, \*\*\**p*<0.001, \*\*\*\**p*<0.0001) (D) CT26 cells with functional miR-424 (WT, miRi-Ctrl, CRISPR-Ctrl, miR-424OE) and CT26 tumor cells without functional miR-424 (miR-4242i, miR-424KO) were used to induce tumor formation in immune competent BALB/c mice. (n=10-20, Kaplan-Meier method for survival curve, log-rank test for survival analysis, \*\*\*\**p*<0.0001) (E) Antitumor specific immune response was detected in tumor free mice that rejected the miR-424i CT26 cells. (Mean±SEM, n=6, t-test, \**p*<0.05, \*\*\**p*<0.001) (F) Depletion of the functional miR-424 did not affect CT26 tumor growth pattern in SCID mice. Proliferation (ki-67) and apoptosis (cleaved-caspase-3) of tumor cells were not influenced by functional miR-424 depletion. (Mean±SEM, n=4, t-test, n.s. no significance) (G) Upper panels: Differentially expressed genes in CT26 cells with and without functional miR-424. Lower panel: Gene Ontology analysis of biological pathways. See also Figure S4 and Table S1-2.

Next, we tested the impact of tumor cell derived miR-424 on tumor formation and growth. Notably, when functional miR-424 was depleted in tumor cells and TEVs, tumor formation and growth rates decreased significantly (Figures 4D and S4A). In the host mice that rejected miR-424i tumors (lacking functional miR-424), we examined antitumor specific immune response (Figures 4E and S4B). In TdLNs and spleen of these mice, we found an expansion of gp70 tumor antigen specific CD8^+^ T cells (Figure 4E). gp70 is a known tumor antigen present in CT26 cells. We got a similar result in the MC38 tumor model with p15E tumor antigen specific tetramer staining (Figure S4B). These data indicated that a robust antitumor immune response could contribute to the rejection of tumor formation in those mice. miRNAs have multiple autonomous targets in cell function.

Therefore, to confirm that the antitumor immunity was the predominant reason for the slower development of tumors lacking functional miR-424, we repeated the experiments in severe combined immunodeficient (SCID) mice, *Cd28*^*-/-*^ mice, and *Cd80/86*^*-/-*^ mice (Figures 4F and S4A). Tumor growth rate, proliferation (Ki-67^+^), and apoptotic cell (cleaved-caspase-3^+^) frequency were similar among tumors with/without functional miR-424 in SCID mice (Figure 4F). Blocking miR-424 did not alter tumor growth in *Cd28*^*-/-*^ and *Cd80/86*^*-/-*^ mice (Figure S4A). Transcriptome analyses of tumor cells revealed that blocking functional miR-424 did not extensively regulate cell-autonomous gene expression in tumor cells (Figures 4G and S4C-E). Taken together, these results indicated that blocking tumor cell-derived miR-424 effectively inhibits tumor development in a CD28-CD80/86 dependent manner.

### Blocking tumor cell derived miR-424 enhanced adaptive antitumor immunity and sensitizes advanced tumors to ICBT

Since blocking tumor cell derived miR-424 suppressed tumor growth by an immunity dependent manner, we then analyzed the impacts of tumor cell derived miR-4242 on anti-tumor immune effectors. Immunohistological analyses on the CT26 and MC38 tumor sections showed that decreasing tumor cell-derived miR-424 led to a higher T-cell infiltration (Figures 5A and S5A). T cells were observed in both peri- and intra-tumor areas (Figures 5A and S5A). Analyses of antitumor immune landscapes were performed in tumor tissues, TdLNs, and the spleen of tumor-bearing mice (Figures 5B-C and S5B-D). Blocking tumor cell-derived miR-424 predominantly altered the antitumor immune landscape in both CT26 and MC38 tumor microenvironments (Figures 5B and S5B) but not in the TdLNs and spleen of tumor-bearing mice (Figures 5C and S5C-D). We found that T cells in tumors that lack miR-424 had elevated CD28 expression compared to T cells in control tumors. Similarly, the CD80 expression on DCs in miR-424 blocked tumors was higher than the control tumors (Figures 5B and S5B). A higher concentration of IL-2 was observed in miR-424 blocked tumors as well (Figures 5B and S5B). Other parameters such as total immune infiltration, T-cell infiltration and proliferation were enhanced by blocking tumor cell-derived miR-424 (Figures 5B and S5B). Remarkably, the major immunosuppressive mechanisms such as tumor tissue PD-L1 expression and PD-1 expression on CD4^+^ T cells were higher in the miR-424 blocked tumors (Figures 5B and S5B). The frequencies of Treg and myeloid-derived suppressor cells (MDSCs) showed a trend of increase (Figures 5B and S5B). These mechanisms indicate a potential negative feedback response.

**Figure 5.**
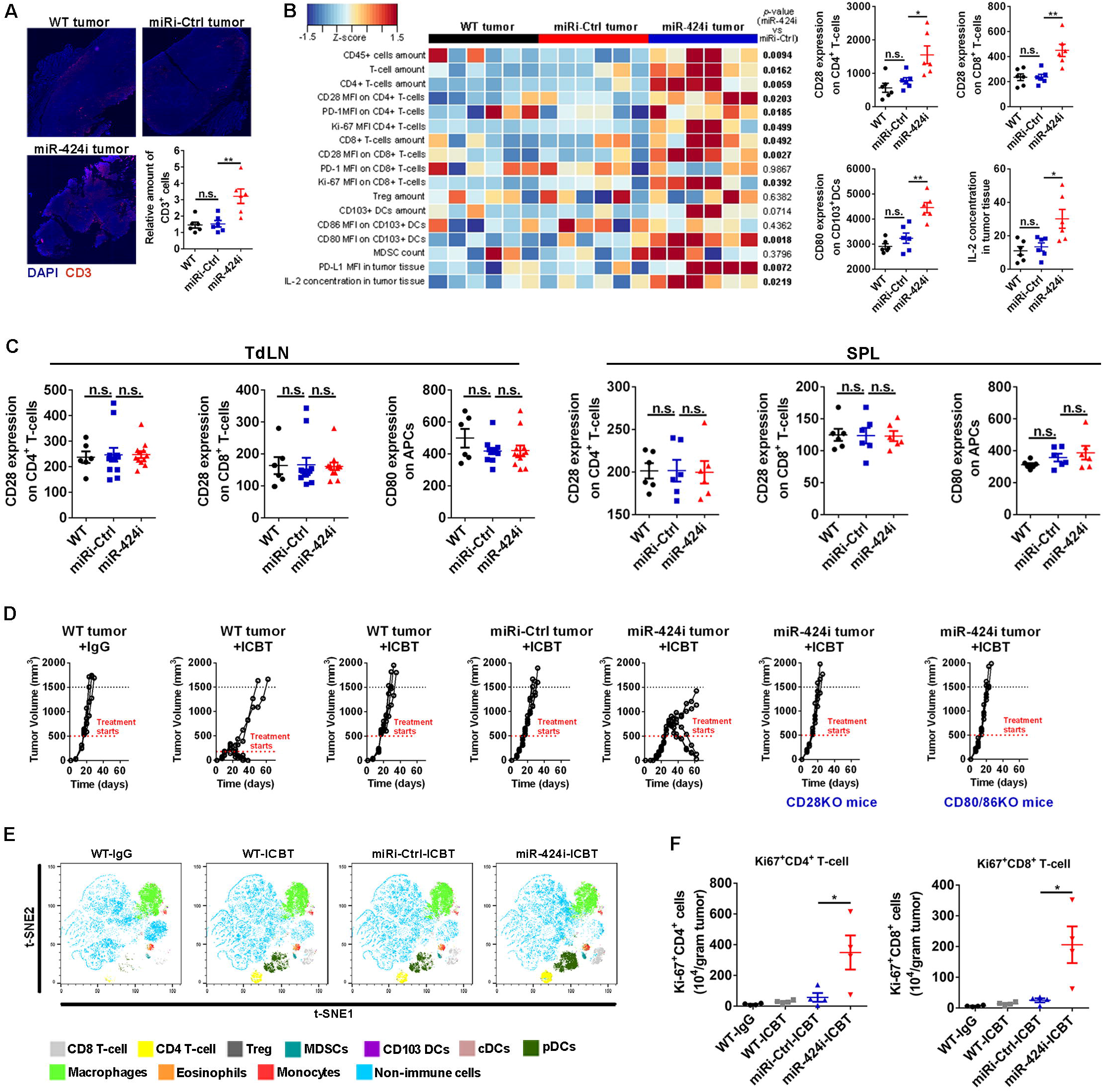
Depletion of functional miR-424 enhanced the costimulatory molecules CD28 and CD80 expression, antitumor immunity, and response to ICBT. (A) The overall T-cell infiltration in CT26 tumors with (WT, miRi-Ctrl) and without (miR-4242i) functional miR-424. T cells were observed in both peripheral and central region of tumors. (Mean±SEM, n=6, t-test, \*\**p*<0.01 n.s. no significance) (B) Heatmap representation of antitumor immune profiles in CT26 tumors without functional miR-424 and control tumors. Right panels: Levels of CD28 and CD80 expression on tumor infiltrating CD4^+^ and CD8^+^ T cells and CD103^+^ DCs. Intratumoral IL-2 concentration were measured in all three groups. (Mean±SEM, n=6, t-test, \**p*<0.05, \*\**p*<0.01 n.s. no significance) (C) Expression of CD28 on T cells and CD80 on antigen presenting cells (APCs) in tumor draining lymph nodes (TdLNs) and spleen (SPL) of MC38 tumor bearing mice. (Mean±SEM, n=6, t-test, n.s. no significance) (D) In MC38 tumor model, ICBT is effective for treating the early stage tumors (volume ∼100mm^3^) but not for controlling the large tumors (volume 500-700mm^3^). Blocking functional miR-424 in tumor cells enhanced the response of established tumors to ICBT in WT C57BL/6 mice, but not in *Cd28*^*-/-*^ mice and *Cd80/86*^*-/-*^ mice. (E) Mass cytometry analysis of immune landscape in established tumors treated with ICBT. (n= 4 cases/condition). A representative t-SNE plots showing immune cell infiltration for each condition. (F) Tumor infiltrating T cells proliferation induced by ICBT in established tumors without functional miR-424. (Mean±SEM, n=4, t-test, \**p*<0.05) See also Figure S5.

Our observations point out that the expression of CD28 on T cells and CD80 on DCs was downregulated in advanced tumors with elevated miR-424 (Figures 2G and Figure S2C). Because CD28 and CD80 are directly involved in the efficacy of anti-PD-1/PD-L1 treatments(Hui et al., 2017; Kamphorst et al., 2017; Sugiura et al., 2019; Zhao et al., 2019), we reasoned that blocking tumor cell-derived functional miR-424 would enhance tumor response to ICBT. We tested anti-PD-1/CTLA-4 treatments in early stage tumors and established tumors with/without functional miR-424 expression (Figures 5D and 5SE). Notably, anti-PD-1/CTLA-4 treatments in established WT tumors were less effective than in early stage tumors (Figures 5D and 5SE). In contrast, blocking tumor cell-derived miR-424 in combination with ICBT caused tumor rejection or reduction in established tumor growth in WT mice. However, the treatment is ineffective in *Cd28*^*-/-*^ mice and *Cd80/86*^*-/-*^ mice even with blocked miR-424 in tumor cells (Figures 5D and 5SE). Systemic assessments by mass cytometry demonstrated distinctive antitumor immune landscapes in anti-PD-1/CTLA-4 treated established tumors with/without miR-424 (Figure 5E). A favorable phenotype with high tumor infiltrating T-cell frequency and function was only seen in treated tumors that lacked functional miR-424 (Figure 5E-F). These data support the significance of tumor cell-derived miR-424 in determining tumor response to ICBT.

### Depleting functional miR-424 induces antitumor immunogenicity of TEVs

Wild type TEVs are generally considered immunosuppressive(Daassi et al., 2020; Whiteside, 2016). On the other hand, TEVs do contain tumor antigens that can induce an antitumor immune response(Diamond et al., 2018; Horrevorts et al., 2019; Poggio et al., 2019; Wolfers et al., 2001). Therefore, we speculated that depleting immunosuppressive factors, such as miR-424, would potentially enhance the immunogenicity of TEVs. To address this, we systemically administered TEVs with/without functional miR-424 to naïve mice. We first validated that the systemically administered TEVs could be taken in by immune cells in the peripheral lymphatic organs (Figure 6A). We then found that administration of TEVs lacking miR-424 (termed modified TEVs) could stimulate the expansion of tumor antigen specific CD8^+^ T cells in peripheral lymphatic organs of naïve mice without inducing extensive non-specific T-cell activation (Figure 6B). Moreover, to investigate whether a modified TEVs induced tumor specific immune response can reject tumors *in vivo*, we induced tumors in mice preconditioned with TEVs (Figure 6C). All mice treated with saline and TEVs with functional miR-424 developed CT26/MC38 tumors (Figure 6C). Remarkably, mice treated with modified TEVs were immunized to the tumor cells or only developed minimal CT26/MC38 tumors (Figure 6C). Repetition in *Tlr4*^*-/-*^ mice showed similar results, indicating that the antitumor specific immune response, rather than potential non-specific immune responses, accounts for the protective effects of modified TEVs (Figure 6C). In SCID mice, *Cd28*^*-/-*^ mice, and *Cd80/86*^*-/-*^ mice, the treatment failed to prevent tumor formation (Figure 6C), indicating that the effect is highly dependent on the CD28-CD80/86 costimulatory pathway.

**Figure 6.**
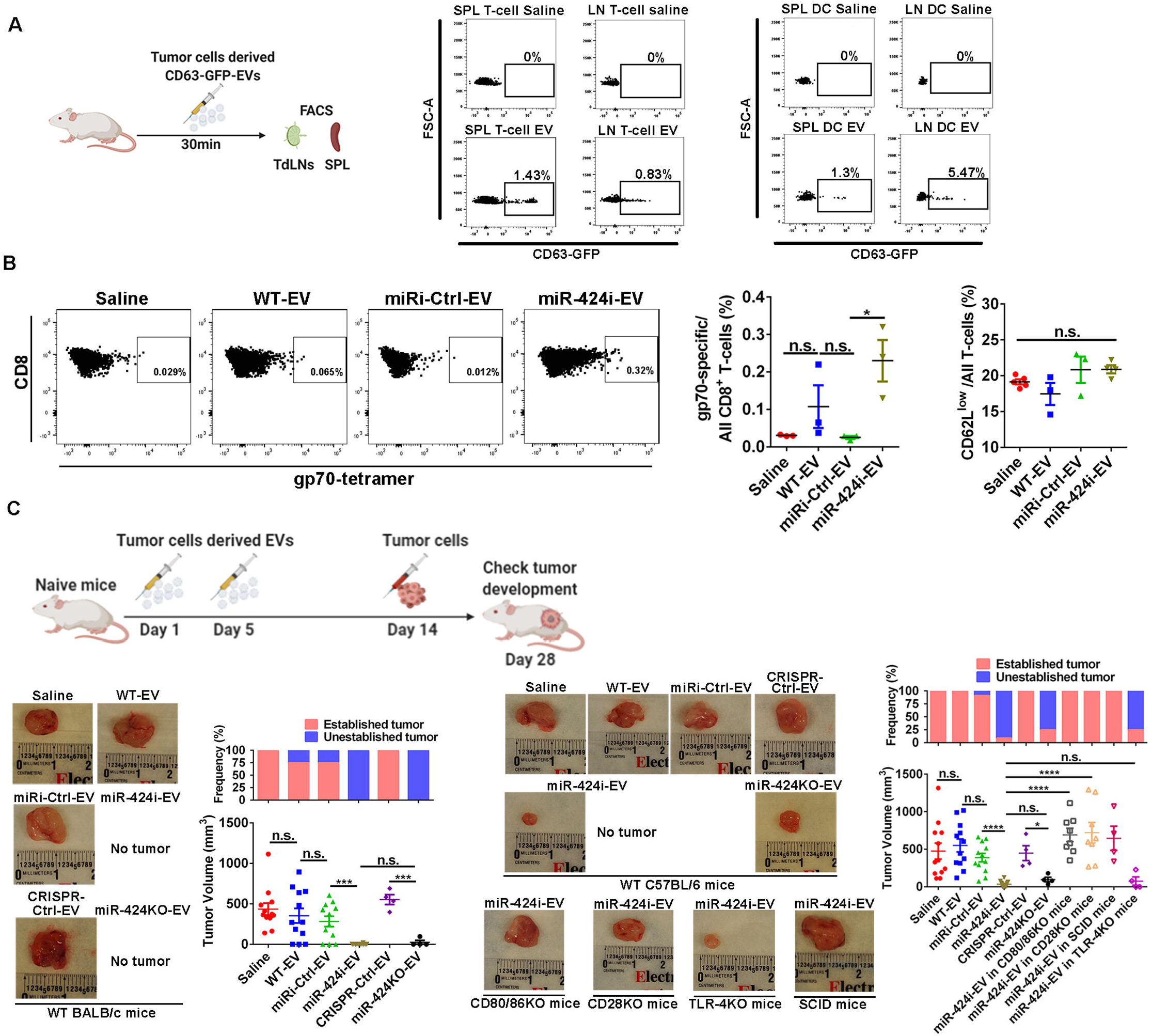
TEVs without functional miR-424 are efficient in stimulating antitumor immune response. (A) Representative flow cytometry dot plots showing intravenously injected TEVs (CD63-GFP labeled) were taken by T cells and DCs in the peripheral lymphatic organs (n=3/group). (B) Tail vein injection of TEVs (10µg) with different status of miR-424 function in naïve mice. TEVs without functional miR-424 (miR-424i-EV) stimulate tumor antigen specific (gp70) T-cell expansion in the peripheral lymphatic organs. Non-specific T-cell activation (CD62L^-^ frequency) was also evaluated. (Mean±SEM, n=4, t-test, \**p*<0.05 n.s. no significance) (C) TEVs (10µg) with different miR-424 functional status was injected to naïve mice intravenously. Two weeks after injection, the mice were challenged with syngeneic tumor cells subcutaneously. Naïve WT mice preconditioned by TEVs without functional miR-424 (miR-424i-EV and miR-424KO-EV) showed lower tumor burden than mice preconditioned by TEVs with functional miR-424 or saline. The experiments were repeated in *Tlr4*^*-/-*^ mice, SCID mice, *Cd28*^*-/-*^ mice, and *Cd80/86*^*-/-*^ mice. The tumor burden was measured two weeks after tumor cells inoculation. Left half: CT26 model; Right half: MC38 model. (Mean±SEM, n=4-12, t-test, \**p*<0.05, \*\*\**p*<0.001, \*\*\*\**p*<0.0001, n.s. no significance) See also Figure S6.

We monitored body weight, blood thrombotic status, and serum cytokine levels of recipient mice (Figure S6A-C) to evaluate treatment safety. No differences were observed between untreated and treated groups (Figure S6A-C). Histological analysis of the major organs was performed and confirmed that the short-term modified TEV treatment (2 doses in one week, i.v., 10μg protein per dose) did not cause severe damage to the heart, liver, lung, kidney, small intestine, or spleen (Figure S6D). Cumulatively, our findings highlighted that modified TEVs are immunogenic and can stimulate antitumor immunity in a safe manner.

### TEVs with miR-424 knockdown enhance ICBT efficacy in an advanced CRC pre-clinical tumor model

Given the potent antitumor immunogenic features of the modified TEVs, we further examined their therapeutic value in advanced CRC pre-clinical tumors. As a preliminary test, we administered the modified TEVs as a treatment for early (100-200mm^3^) and established (500-700mm^3^) subcutaneous tumors and found that the modified TEVs treatment alone was effective in minimal tumors, but not advanced tumors (Figure S7A). Since we have shown that the advanced tumors have high levels of immunosuppressive factors (Figure 2G), we reasoned that the antitumor immune response initiated by modified TEVs was not sustained in the advanced tumors. Thus, we proposed to combine the modified TEVs with ICBT to treat immunosuppressive tumors.

To mimic human late-stage CRC, we developed a cecum based orthotopic transplantation pre-clinical model(Cespedes et al., 2007; Song et al., 2018; Terracina et al., 2015). Both established primary tumor in cecum and metastatic lesions in the liver and peritoneal cavity were observed in this model (Figure S7B-C), indicating the advantages of using it to study treatment for stage IV CRC. For established tumors, two doses of modified TEVs were given during the 1^st^ week of treatment for induction and anti-PD-1/CLTA-4 were given twice a week until the endpoint for sustention (Figure 7A). Notably, combination of modified TEVs with ICBT significantly reduced tumor metastatic burden and extended overall survival to more than 50% compared with ICBT only group (Figure 7B-C). Tumors in each group were subjected to immune cell analyses which showed that the combination treatment with modified TEVs and ICBT stimulated the most potent antitumor immunity with the highest T-cell infiltration frequency and activity (Figures 7D and S7D). Thus, combination treatment using modified TEVs that can stimulate tumor specific T-cell expansion in peripheral lymphatic organs along with ICBT that depletes immunosuppressive factors in the TME, highlights a potentially novel therapeutic approach to delay progression in late-stage CRC.

**Figure 7.**
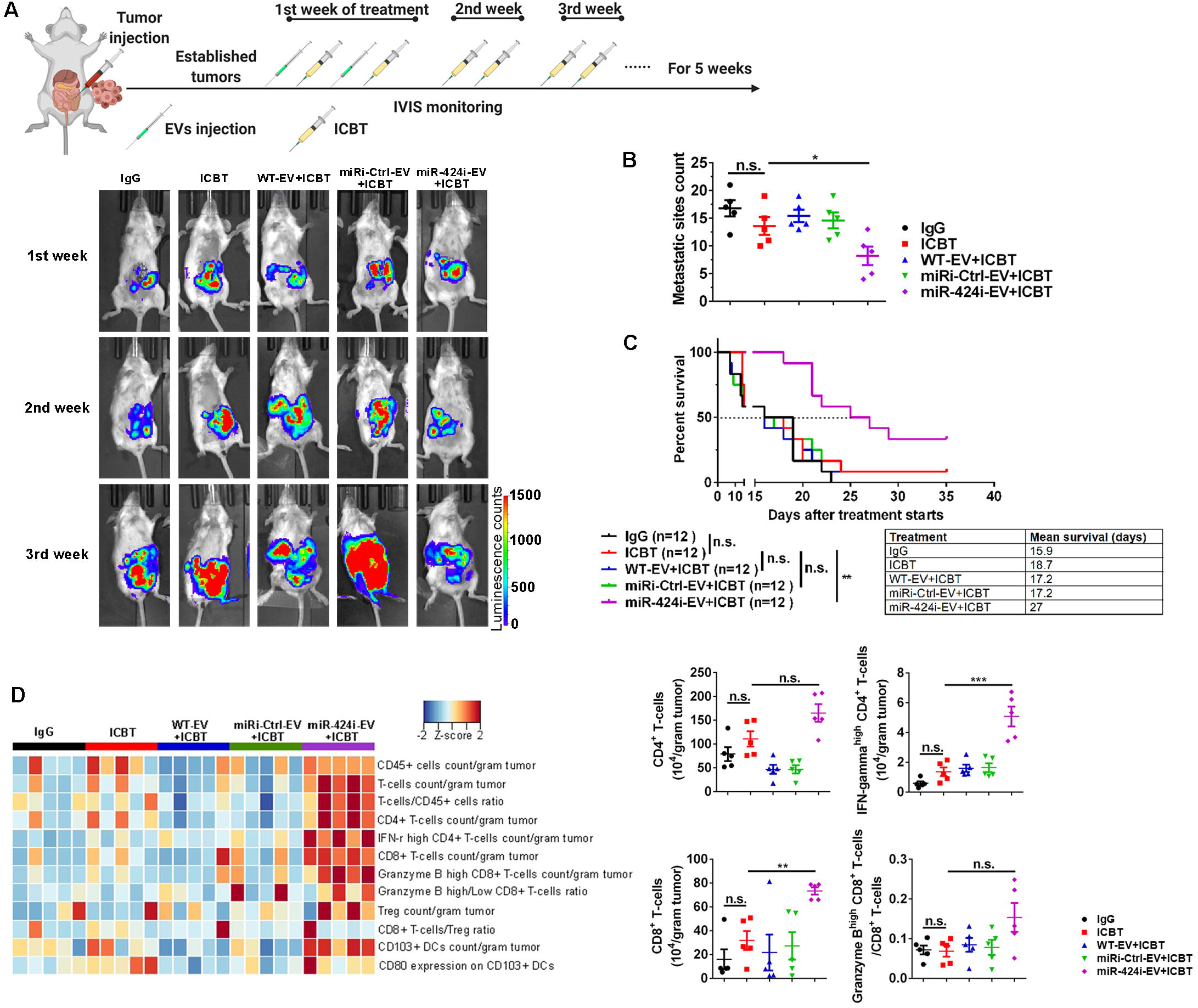
TEVs without miR-424 enhanced the efficacy of ICBT in established orthoptic tumors. (A) The schematic of treatment design. CT26-Luc cells were implanted to the cecum of naïve mice to establish an advanced stage orthotopic tumors. Tumor development was monitored by periodic *in vivo* imaging. Mice with established tumors were assigned to different treatments (see methods for dose details, representative images from each treatment groups are shown). Mice were either sacrificed after two weeks of treatment for evaluating disease burden and immune infiltration or followed up for survival analysis. (B) Metastatic tumor burden evaluated after two weeks of treatment. (Mean±SEM, n=5, t-test for two group comparisons, one-way ANOVA for multiple groups comparison, \**p*<0.05, n.s. no significance) (C) Kaplan-Meier overall survival analysis of mice in each group. The mean survival time listed in the table. (n=12, log-rank test for survival analysis, \*\**p*<0.01, n.s. no significance) (D) Heatmap representation of immune profiles of tumors treated with different therapies. Right panels show quantitative analysis. (Mean±SEM, n=5, t-test for indicated two groups, \*\**p*<0.01, \*\*\**p*<0.001, n.s. no significance) See also Figure S7.

## DISCUSSION

In this study, we demonstrate a critical immunosuppressive role of tumor cell-derived miR-424 as a negative regulator of CD28-CD80/86 costimulatory signaling in tumor infiltrating T cells and DCs.

Blocking miR-424 in TEVs alleviated the immunosuppressive effects and enhanced the immune stimulatory effects of TEVs, making them a potential adjuvant treatment for ICBT used to treat CRC. It was noteworthy that the CT26 and MC38 cells used in this study are representative of CRC-MSS and -MSI phenotypes, respectively(Efremova et al., 2018). Thus, our data revealed a shared mechanism of T-cell costimulation in these CRC subtypes.

In both men and women, CRC is the third most common type of cancer(Siegel et al., 2020). CRC patients were among the first cohort of cancer patients tested for ICBT; however, the majority of late stage CRC patients were unresponsive to ICBT in the early clinical trials. After several trials, it became evident that almost all responders had the MSI subtype(Brahmer et al., 2010; Le et al., 2015). Although the mutational loads in different CRC subtypes are considered as one of the major determining factors of ICBT response(Zhao et al., 2018), other mechanisms regulating immune response are likely to exist. Indeed, around 40-70% of the immune infiltrated late stage MSI CRC with high mutational load failed in ICBT(Le et al., 2015; Oliveira et al., 2019). Furthermore, MSS CRC that are considered resistant to ICBT can still have a high frequency of tumor infiltrating T cells(Kikuchi et al., 2019; Mlecnik et al., 2016; Pages et al., 2018). Our observation in TCGA CRC dataset and a cohort of 71 CRC patient samples validated that a proportion of the MSS tumors have comparable T-cell infiltration as with MSI tumors (Figure 1). This evidence supports the hypothesis that independent of the MSI/MSS classification, there are other potential mechanisms controlling the adaptive immune response in CRC which needs further investigation.

ICBT functions through blocking the inhibitory immune checkpoint molecules expressed on tumor cells, T cells, and other TME components. However, to rescue exhausted T cells by blocking PD-1/PD-L1 signaling, positive costimulation from the CD28-CD80/86 pathway is required(Kamphorst et al., 2017). Furthermore, biochemical studies demonstrated that T-cell expressed CD28 is a primary target for PD-1 mediated inhibition(Hui et al., 2017). In infectious conditions, CD28 expression is also required after T-cell priming for helper T-cell-mediated immune protection(Linterman et al., 2014) and the presence of CD28-mediated signaling prevents induction of anergy in T cells(Harding et al., 1992). All of these observations prompted us to investigate whether the CD28-CD80/86 costimulatory signaling is a prerequisite for ICBT response and whether this signaling is preserved in the CRC immune microenvironment. Remarkably, our analysis of human CRC indicated that some tumor infiltrating T cells and DCs lack expression of CD28 and CD80/86, respectively (Figure 1). Moreover, depleting or blocking the CD28-CD80/86 costimulatory signaling nullified the efficacy of ICBT in mouse CRC models that have immune cell infiltrations (Figure 1). Our findings are consistent with early reports in non-small cell lung cancer and melanoma tumors(Kamphorst et al., 2017; Li et al., 2010) and support the hypothesis that the lack of intact CD28-CD80/86 costimulatory signals on infiltrated T cells and DCs restricts ICBT response in human CRC.

In non-cancerous conditions, CD28 expression levels decrease during antigenic exposure but rapidly revert back to pre-stimulation levels. However, sustained T-cell stimulation causes CD28 downregulation and eventual loss. CD28 downregulation with T-cell activation predominantly involves transcriptional repression and increased protein turnover as a T-cell intrinsic negative feedback mechanism(Weng et al., 2009). Biosynthesis and expression of CD80/86 on DCs is stimulated by antigen presentation and is associated with DC maturation. In cancerous conditions, the effects of tumor and stromal cells on T cells and DCs through direct contact and/or paracrine mechanisms cannot be ignored(Xie et al., 2019; Zhao and Subramanian, 2017, 2018). Our previous studies and other reports have established that abnormal expression of miRNAs is associated with CRC pathobiology(Li et al., 2014; Moridikia et al., 2018; Sarver et al., 2009).

Therefore, we sought to investigate tumor cell-secreted miRNAs as a tumor intrinsic mechanism of CD28-CD80/86 downregulation in immune cells. We identified that miR-424s negatively regulate CD28 and CD80 expression (Figure 2). miR-424 was upregulated in CRC compared to normal colon, T cells, and DCs (Figures 2 and 3). In tumors, miR-424 was upregulated in tumor cells and was transported to surrounding T cells and DCs via TEVs, leading to downregulation of CD28 and CD80 proteins and defective T-cell function (Figure 3). Increased miR-424 levels in tumor cells were induced by hypoxia and accompanied by immune exclusion (Figure 2 and 3). These data validated that tumor intrinsic mechanisms can be an initiating factor for immunosuppression. Of course, we cannot exclude other unknown mechanisms that upregulate miR-424. The immune editing during tumor development may also contribute to the selection and accumulation of miR-424 producing tumor cells.

In mouse tumors, depleting or blocking miR-424 in tumor cells significantly delayed tumor growth in an immune-dependent manner (Figure 4). Immune landscape analyses demonstrated that lack of tumor derived miR-424 rescued CD28 expression on both tumor infiltrating CD4^+^ and CD8^+^ T cells. CD80 expression on the tumor infiltrating CD103^+^ DCs that are critical for tumor antigen presentation and effector T-cell trafficking(de Mingo Pulido et al., 2018; Roberts et al., 2016; Salmon et al., 2016; Spranger et al., 2017) were increased in tumors lacking miR-424. Notably, T-cell frequency and function were significantly higher in those tumors. As a consequence, lack of tumor-derived miR-424 and restoration of CD28-CD80/86 signaling reversed tumor intrinsic resistance to anti-PD-1/CTLA-4 treatment (Figure 5). Our data reinforces previous findings that intact CD28-CD80/86 costimulatory pathways could attenuate PD-1 mediated T-cell exhaustion and were associated with better response to anti-PD-1 treatment(Fenwick et al., 2019; Kamphorst et al., 2017; Mizuno et al., 2019). Recent mechanistic studies describing the immunostimulatory and therapeutic significances of CD80 expression on APCs(Sugiura et al., 2019; Zhao et al., 2019) also support our findings. In summary, our studies establish that TEVs delivered miR-424 is a crucial mechanism causing defective CD28-CD80/86 costimulatory signaling in human CRC that can induce ICBT resistance.

Blocking tumor cell derived miR-424 restored the defective CD28-CD80/86 costimulatory signaling in tumors and resulted in a better response to ICBT (Figure 5). However, in clinical conditions, it is difficult to genetically alter tumor cells or specifically block miRNAs in the TME. TEVs carry tumor antigens and are being tested as potential cell-free cancer vaccines(Diamond et al., 2018; Wolfers et al., 2001). Therefore, to translate our mechanistic findings to potential therapeutic approaches, we considered enhancing the immunostimulatory aspects of TEVs by eliminating the miR-424 induced immunosuppression. Our modified TEVs (without immune suppressive miR-424) stimulated a tumor-specific immune response that was independent of the tumor’s endogenous antigen presentation and T-cell priming. Systemic administration of modified TEVs in naïve mice induced tumor-specific immune responses and are protective of tumor cell challenge in the preconditioned mice (Figure 6). Careful monitoring of the health of the recipient mice indicated there were no serious adverse side effects of this treatment (Figure S6). A recent phase 2 stage clinical trial suggested that utilizing the anti-PD-1/anti-CTLA-4 combination treatment as a neoadjuvant therapy in early stage CRC might have a better response than being used as an adjuvant therapy in late stage tumors. Therefore, we established an orthotopic CRC model that mimics human metastatic CRC tumors to test the therapeutic efficacy of modified TEVs(Chalabi et al., 2020). In such an aggressively progressing CRC model, we successfully tested combining the modified TEVs with ICBT (Figure 7). This combination resulted in an increased immune response and mouse survival rate over the treatment with ICBT alone (Figure 7). This is most likely due to the modified TEVs ability to stimulate anti-tumor specific T-cell responses and that could be sustained by ICBT. Similarly, a recent study using TEVs produced by irradiated mouse breast cancer cells demonstrated the enhanced costimulatory functions of DCs(Diamond et al., 2018). These findings together with our results underscore the potential of using modified TEVs with enhanced immunostimulatory effects in stimulating antitumor immunity and treating ICBT resistant tumors.

Our results have several important implications for clinical translation. First, we show that tumor cells may induce positive costimulatory pathways deficiency in tumor infiltrating immune cells. To revitalize the tumor infiltrating T cells, protecting or restoring the positive costimulatory pathways may be equally important as suppressing the inhibitory pathways. This concept is backed by a recent report using CD28 stimulatory mechanisms to synergize with anti-PD-1 blocking antibody(Fenwick et al., 2019). Second, our data showing that modified TEVs could stimulate anti-tumor T-cell responses and reverse ICBT resistance in advanced CRC models suggest potential therapeutics for ICBT resistant tumors. The WT TEVs are considered as immunosuppressive in general(Iero et al., 2008; Kurywchak et al., 2018). However, recent findings identified the immunosuppressive factors in those TEVs and showed that blocking them can alleviate the immunosuppression(Chen et al., 2018; Kim et al., 2019; Poggio et al., 2019). Further, adding immunostimulatory factors into the TEVs can even enhance their ability to initiate immune responses(Diamond et al., 2018). Traditional TEVs related therapies used WT TEVs to pulse DCs *in vitro* and TEVs loaded DCs were infused to treat tumors(Bu et al., 2011; Gu et al., 2015; Zuo et al., 2020). This method avoids the potential negative effects of TEVs on the immune system *in vivo*. However, this approach is time consuming and did not consider the negative effects of WT TEVs that may impart functional defects in DCs, leading to treatment failure. Administration of the modified TEVs can minimize the negative effects of WT TEVs on immune response and, at the same time, efficiently stimulate tumor-specific immunity *in vivo*. Modified TEVs could be an alternative to the current TEVs based cancer treatments.

Taken together, our study reinforces previous reports regarding positive costimulatory pathway and TEVs in ICBT(Kamphorst et al., 2017; Poggio et al., 2019). We deciphered a novel mechanism of the CRC immune evasion and an innovative concept of modified TEVs based treatment. It is noteworthy that miR-424 may have strong effects on other immune cell populations or immune processes considering it has a large number of predicted immune gene targets. Our current results do not exclude the contribution of other potential immune evasive mechanisms of ICBT failure(Kalbasi and Ribas, 2020). A comprehensive understanding of immune suppressive mechanisms and accurate blocking of these suppressive factors is critical and necessary for a successful treatment. Finally, an array of potent immunosuppressive factors have been identified as well(Kim et al., 2019; Poggio et al., 2019; Ricklefs et al., 2018; Vignard et al., 2020). Blocking a bulk of these negative regulators may further enhance the efficacy of modified TEVs on stimulating anti-tumor immunity but requires more investigations.

## ACKNOWLEDGMENTS

This work is supported by the Minnesota Colorectal Cancer Research Funds, Mezin-Koats Colon Cancer Research Award and the Department of Surgery, Research funds. We thank Dr. Lihua Li for generating the preliminary results. We also thank Mr. Nile Liu, Ms. Duha Al Sherriff, Audrey McCarthy, and Beminet Kassaye for technical assistance. We thank the comparative oncology core facility and Mass Cytometry core facility at University of Minnesota for helpful assistance. We also thanks Drs. Andrew Nelson and Aqsa Nasir for help with CRC tissue section and IHC analysis. We thank Drs. Timothy Starr, Christopher Pennell, Emil Lou and Kaylee Schwertfeger for critical comments and feedback on the manuscript. We thank Drs. Marc Jenkins and Kris Hogquist for providing CD28 and CD80 knockout mouse and helpful comments.

## AUTHOR CONTRIBUTIONS

X.Z. and S.S. conceived and designed the experiments. X.Z. and D.W. performed all experiments and data analyses. C.Y., and D.W. provided assistance in gene expression analysis and tumor organoid culture. X.Z., C.Y., D.W. and S.S. commented on the data. X.Z. and S.S. co-wrote the paper. S.S. supervised this project.

## DECLARATION OF INTERESTS

The authors declare no competing interests

## Sequence data

**RNA sequence accession number**

## Notes

### Competing Interest Statement

The authors have declared no competing interest.

